# When is spontaneous formation of spatial patterns robust?

**DOI:** 10.1101/2024.10.30.621030

**Authors:** Vit Piskovsky, Philip K. Maini

## Abstract

From atomic spins in magnets to galaxies, and from embryonic development to patchy vegetation in arid environments, the physical world is filled with complex systems that spontaneously form spatial patterns. While simple mathematical models, such as reaction-diffusion systems, can explain the formation of such patterns, the complexity of the physical systems these models aim to describe necessitates the analysis of model robustness. Our work utilizes random matrix theory to provide arguably the first definition of robustness that is analytically tractable, providing an easy guide for identifying spatial interactions that robustly generate spatial patterns. We illustrate our theory on examples from mathematical biology, showing that diffusion alone cannot robustly generate spatial Turing patterns in large and unstructured systems, while advection, chemotaxis and non-local interactions can robustly promote pattern formation. Furthermore, we use our theory to prove that the spinodal decompositon of soft condensed matter is dynamically robust and predict the dynamics beyond regimes permitted by the standard Landau-Ginzburg theory. By classifying different spatial interactions based on the robustness of pattern formation, this work provides insights into which mechanisms are fundamental for pattern formation in large and unstructured physical systems.

---

Spatial patterns emerge across all scales of the physical world and their spontaneous formation provides a central theme of scientific study. The spins of atoms self-organize in magnetic materials [1, 2], reacting chemicals spontaneously form patterns [3], bacterial cells create spatially heterogeneous biofilms [4], animals develop patterned skin pigmentation [5], vegetation is patchily distributed in arid environments [6], and planets, solar systems and galaxies spontaneously emerged from a relatively homogeneous bulk of energy and matter after the Big Bang [7]. Even though different mechanisms drive pattern formation in these systems, the spontaneous formation of spatial patterns has a similar mathematical structure.

Specifically, these systems are commonly described by *a* = 1, …, *N* field variables *n*_*a*_(*t*, ***x***) that represent the physical quantities of interest, ranging from magnetization, chemical concentrations, biofilm bacteria or vegetation to the matter density in the Universe, at time *t* and spatial coordinates ***x*** = [*x*_1_, …, *x*_*D*_]^*T*^. The dynamics of these variables are often described by equations of the form

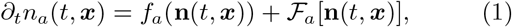

where **n**(*t*, ***x***) = [*n*_1_(*t*, ***x***), …, *n*_*N*_ (*t*, ***x***)] is a vector of field variables, *f*_*a*_ describes temporal interactions and ℱ_*a*_ describes spatial interactions, that is, ℱ_*a*_ is an operator such that

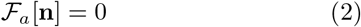

for any spatially homogeneous state **n**(*t*, ***x***) = **n** (Fig. 1a). For example, the condition (2) is satisfied if the domain is infinitely large ***x*** ∈ ℝ^*D*^ and

**FIG. 1.**
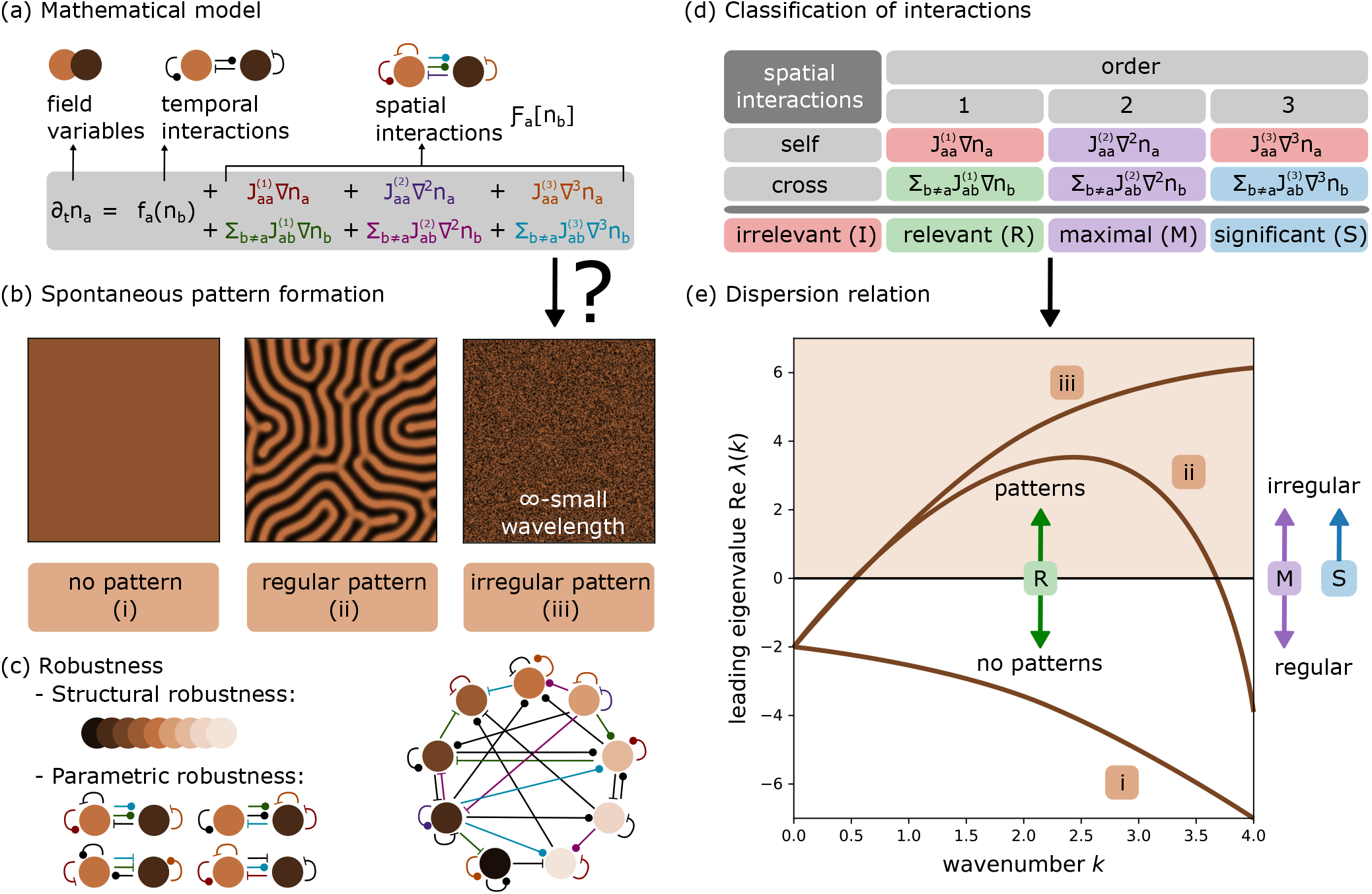
Robustness of spontaneous pattern formation. **(a)** Mathematical model for field variables *n*_*a*_(*t, x*) (brown circles) with temporal interactions ℱ_*a*_(*n*_*b*_) (black connections) and spatial interactions *F*_*a*_[*n*_*b*_] (coloured connections), which are expanded order by order with matrix coefficients 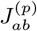. **(b)** Spatial patterns form spontaneously if a temporally stable homogeneous stationary state is unstable to spatial perturbations. Several outcomes are possible: (i) no patterns, (ii) regular patterns (with finite wavelengths), and (iii) irregular patterns (with infinitely small wavelengths). **(c)** A key question is whether patterns form robustly in a given model, that is for different numbers *N* of field variables (structural robustness) and for different parameter values of interactions 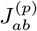 (parametric robustness). We define pattern formation to be robust if patterns can form spontaneously in a system with many variables *N → ∞* and independent random interaction parameters 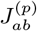. **(d)** To analyze robustness of spatial interactions, different interactions are classified as irrelevant (I, red), relevant (R, green), maximal (M, purple) and significant (S, blue), as explained in Definition 1. **(e)** The dispersion relation captures the growth rate (leading eigenvalue) of perturbations with wavenumber *k* around the homogeneous stationary state. Therefore, patterns form spontaneously only if some perturbation has a positive growth (brown region) and pattern formation is regular if perturbations with infinitely large wavenumbers have negative growth (white region). Different dispersion relations (brown curves) can be identified with different regimes 1-3 of pattern formation from panel (b). Theorems 1 and 2 explain that relevant interactions influence the robustness of pattern formation, while maximal and significant interactions control its regularity. Coloured arrows indicate changes in the dispersion relation induced by changes in the interaction strength *r*_*p*_ of an interaction from a class in panel (d), see Definition 1.

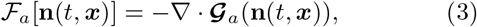

where **𝒢**_*a*_[**n**(*t*, ***x***)] is a flux vector field. In this case, equation (1) is called a continuity equation, which plays a central role in the theory of electromagnetism [8], fluids [9], astrophysical fluids [10], soft condensed matter [11, 12], quantum mechanics [13], thermodynamics [14], chemical reactions [15] and multiple biological systems [16–18]. With the general formulation of the dynamics in equation (1), spontaneous pattern formation can be studied similarly across diverse physical systems as follows.

Without spatial interactions, ℱ_*a*_ = 0, and on infinite domains, ***x*** ∈ ℝ^*D*^, the temporal dynamics in equation (1) commonly admits a stable spatially homogeneous stationary state **n**(*t*, ***x***) = ***n***^*\**^, which is given by

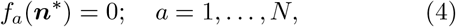

and is associated with a Jacobian matrix 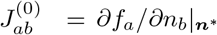 with eigenvalues that all have negative real parts. Importantly, a positive answer to the following question determines spontaneous pattern formation. Is the spatially homogeneous stationary state **n**(*t*, ***x***) = ***n***^*\**^ stable when the spatial interactions ℱ_*a*_ are present (Fig. 1b)?

As can be seen in [2–7], this question has sparked atention across different fields of study. Most notably, spontaneous pattern formation has been studied intensively in the context of reaction-diffusion systems [16], with a long history tracing back to Alan Turing. In 1952, Turing showed that the diffusion,

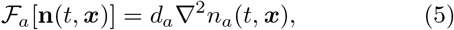

of two interacting variables (*N* = 2) with diffusivities *d*_*a*_ *>* 0 can destabilise their temporally stable spatially homogeneous distribution, even though the diffusion of a single self-interacting variable (*N* = 1) cannot do this [19]. In particular, for patterns to form, one variable must diffuse slowly and activate the dynamics, while the other must diffuse quickly and inhibit the dynamics [16]. Despite the simplicity of the proposed Turing mechanism, observing Turing patterns in experimental and natural systems took several decades [20]. Moreover, those systems that admit Turing patterns are often more complex than can be described with two variables, a slow activator and a fast inhibitor. For example, the chemical reactions that admit Turing patterns involve multiple reactants [3, 21, 22] with similar diffusivities [23]. Biofilm patterns are composed of bacteria that interact with complex chemical environments, secrete multiple extracellular polymeric substances and are capable of biased non-diffusive motility [24]. In zebrafish, pigment cells - melanophores and xanthophores - interact with various chemical factors [5] and undergo a non-diffusive movement [25] to produce stripes and spots on the zebrafish skin. The patterns of vegetation in an arid environment are affected by various biotic and abiotic factors [6], and plant seeds can be dispersed non-locally [26], for example by rodents [27]. The inherent complexity of these systems raises an important question. Is diffusion sufficiently robust to explain pattern formation in such complex physical systems [28]?

In principle, there are different aspects of robustness [28], including parametric and structural robustness (Fig. 1c). Pattern formation is said to be robust parametrically when it occurs for a broad range of parameters in the interactions *f*_*a*_ and ℱ_*a*_, while structural robustness requires pattern formation to occur across a broad range of spatial dimensions *D* and variables *N*. For diffusion, the analytical conditions for pattern formation were recently derived for *N* = 3 variables and necessary conditions were provided for an arbitrary number of variables *N* [29]. However, as these conditions are difficult to interpret, several authors have now investigated the robustness of pattern formation with *N ≥* 2 variables numerically [23, 30, 31]. Inspired by Robert May’s idea to examine the stability of large complex systems with a random matrix theory approach [32], these works accounted for parametric robustness by sampling interactions randomly and for structural robustness by repeating the calculation for multiple numbers of variables *N* [23, 31]. Surprisingly, the probability of observing pattern formation in these random systems rapidly decreased with the increasing number of variables *N*, as has been recently supported by a heuristic analytical argument [33]. In parallel, pattern formation has been analyzed beyond the contexts of diffusion, such as for advection [34], chemotaxis [35], cross-diffusion [36], or non-local interactions [37, 38]. Interestingly, these works often report that such alternative mechanisms promote pattern formation, but the comparison of pattern formation between models with different spatial interactions ℱ_*a*_ is hindered by the absence of a formal and analytically tractable definition of robustness.

In this Letter, we propose arguably the first analytical and systematic approach to quantifying the robustness of pattern formation governed by different spatial interactions ℱ_*a*_. Inspired by random matrix theory, we define pattern formation to be robust if patterns form spontaneously in the considered system when it is large (i.e., includes many variables *N → ∞*) and unstructured (i.e., has random, independent and identically distributed interactions), allowing us to capture both parametric and structural robustness. Utilizing our new definition and the recent progress in random matrix theory [39, 40], we will provide a simple rule to identify which spatial interactions ℱ_*a*_ are congruent with robust formation of spontaneous patterns (Fig. 1d-e).

To formalize this idea, we use linear stability analysis to determine whether spatial perturbations around a temporally stable homogeneous stationary state ***n***^*\**^ can grow, allowing for spontaneous pattern formation [16]. In particular, linearization of equation (1) around the homogeneous stationary state ***n***^*\**^ and a subsequent Fourier transformation show that whether or not the spatial perturbations grow on an infinitely large domain ***x*** ∈ ℝ^*D*^ is determined by the eigenvalues of the matrix

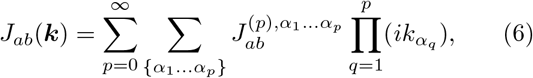

where ***k*** = (*k*_1_, …, *k*_*D*_) is a wavenumber vector of the perturbation,

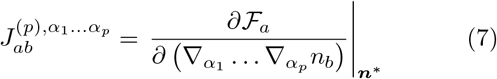

describes different spatial interactions of order *p ≥* 1, and 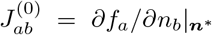 describes the stabilizing temporal interactions and has all eigenvalues with negative real parts. Denoting *λ*(***k***) to be the eigenvalue of *J*_*ab*_(***k***) with maximal real part, it can be noticed that patterns form spontaneously precisely if there exists a perturbation with wavenumber ***k***^*\**^ that can grow at a positive rate

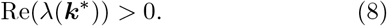

Moreover, the formation of such patterns is regular if perturbations with infinitely small wavelengths cannot grow, that is if Re(*λ*(***k***)) is bounded from above by zero as |***k***| *→ ∞* (Fig. 1e).

Having identified when patterns form spontaneously and regularly, we can return to our definition of robustness. For simplicity of presentation, we will restrict our study to a single spatial dimension (*D* = 1), which allows us to ignore different spatial indices *α*_*q*_ and replace a wavenumber vector ***k*** by a single wavenumber *k*. With the goal to study unstructured systems, we assume that the interactions 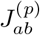 of order *p* between variables *a* and *b* are sampled randomly and independently for different orders *p* and different combinations of *a* and *b*, with self-interactions (*a* = *b*) and cross-interactions (*a* ≠ *b*) of the same order *p* sampled from identical probability distributions, with means and variances given by

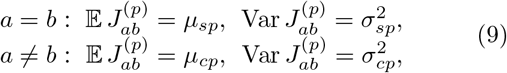

where we account for the possibility that the cross-interactions are symmetric (*τ*_*p*_ = +1), neutral (*τ*_*p*_ = 0) or anti-symmetric (*τ*_*p*_ = −1) via [41]

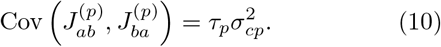

With this assumption, we can say that an interaction is present in the system if the corresponding mean or variance are non-vanishing. This allows us to classify the interactions as follows (Fig. 1d).

## Definition 1

*Let P be the maximal even order of interactions that are present in the system. Then, we classify the spatial interactions (p ≥* 1*) as*

- *maximal (M): interactions of order P*
- *irrelevant (I): symmetric cross-interactions and self-interactions of any odd order p*,
- *significant (S): cross-interactions of odd order p > P that are not symmetric*,
- *relevant (R): all remaining interactions of order p* < *P*.

*For example, if no odd cross-interactions are symmetric, we have:*

| Order | 1 | 2 | 3 | ... | $P-1$ | $P$ | $P+1$ | $P+2$ | $P+3$ | ... |
| --- | --- | --- | --- | --- | --- | --- | --- | --- | --- | --- |
| Self | $I$ | $R$ | $I$ | ... | $I$ | $M$ | $I$ | ∅ | $I$ | ... |
| Cross | $R$ | $R$ | $R$ | ... | $R$ | $M$ | $S$ | ∅ | $S$ | ... |

*Moreover, we define the strength r*_*p*_ *of interactions of order p ≥* 0 *by*

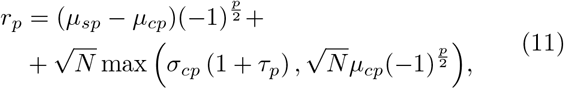

*when p is even, and by*

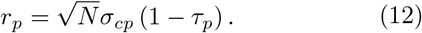

*when p is odd*.

The interaction strengths can be used to simplify the subsequent analysis. As shown in Appendix A, the interaction strengths measure whether the matrix 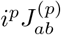 has all eigenvalues with negative real parts (*r*_*p*_ < 0) or not (*r*_*p*_ *>* 0). For example, the assumption about the stability of the homogeneous stationary state ***n***^*\**^ translates to the assumption that *r*_0_ < 0.

Having introduced unstructured systems as those with random interactions, we can now formulate our definition of robustness (Fig. 1c).

## Definition 2

*We say that spontaneous pattern formation is robust in a system given by equation* (1) *if the patterns can form spontaneously when the system is large (N → ∞) and unstructured (i*.*e*., *has random, independent and identically distributed interactions as in equations* (9) *and* (10)).

With these definitions, we can now state the main result (Fig. 1e, see Appendix A for proof).

## Theorem 1

*Assuming that the variance of the self-interactions is bounded by the variance of cross-interactions (i*.*e*., *σ*_*sp*_ *≤ Cσ*_*cp*_ *for some constant C >* 0*), it is true that:*

1. *When irrelevant interactions are the only interactions present, pattern formation is not robust*.
2. *When at least one significant interaction is present with a non-vanishing strength (r*_*p*_ ≠ 0*), pattern formation is robust and irregular*.
3. *When all significant interactions have vanishing strength (r*_*p*_ = 0*) and the relevant interactions are absent, pattern formation is either not robust (r*_*P*_ *≤* 0*) or it is robust and irregular (r*_*P*_ *>* 0*), depending on the strength of the maximal interaction r*_*P*_.
4. *When a relevant interaction is present and sufficiently strong (r*_*p*_ *≫* 0*), pattern formation is robust. Moreover, such pattern formation is regular, provided all significant interactions have vanishing strength (r*_*q*_ = 0*) and the maximal interaction has a negative strength (r*_*P*_ < 0).

Importantly, Theorem 1 allows us to determine whether a particular spatial interaction ℱ_*a*_ in equation (1) forms spontaneous patterns robustly based on a simple inspection of the terms in ℱ_*a*_. For example, if ℱ_*a*_ includes only irrelevant interactions, then Theorem 1 predicts that patterns do not form robustly. One could think that this is a consequence of the additional assumption (*σ*_*sp*_ *≤ Cσ*_*cp*_ for some constant *C*) that restricts the heterogeneity of self-interactions to zero when the cross-interactions are absent. While this additional assumption provides a key simplification in the proof of Theorem 1 (see Appendix A), the key propositions of Theorem 1 remain valid even if all cross-interactions are absent (see Appendix B for proof).

## Theorem 2

*Assume that the temporal interactions are neutral (τ*_0_, *µ*_*c*0_ = 0*) and that the only spatial interactions present in the system are self-interactions (with arbitrary distribution). Then, the following statements are true:*

1. *When irrelevant interactions are the only interactions present, pattern formation is not robust*.
2. *When the relevant interactions are absent, pattern formation is either not robust* 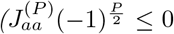 *for all a) or it is robust and irregular (* 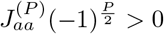 *for some a), depending on the maximal interactions* 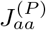.
3. *When a relevant interaction is sufficiently strong (*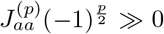 *for some a), pattern formation is robust. Moreover, such pattern formation is regular, provided the maximal interactions are negative* (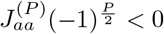 *for all a)*.

Moreover, it is possible to extend these general results to multiple spatial dimensions, *D >* 1, as outlined in Appendix C.

In the rest of this Letter, we will illustrate the general results on several examples from mathematical biology and soft condensed matter physics. In mathematical biology, the spatial interactions of advection, diffusion and chemotaxis play a key role [16]. With these interactions, the dynamics in equation (1) becomes

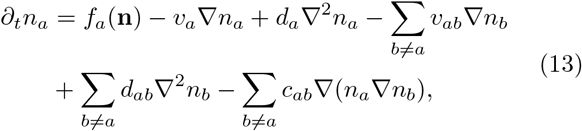

where *v*_*a*_ (resp. *v*_*ab*_) is the velocity of self-advection (resp. cross-advection), *d*_*a*_ (resp. *d*_*ab*_) is the diffusivity of selfdiffusion (resp. cross-diffusion) and *c*_*ab*_ describes chemotaxis of variable *a* up the gradient of variable *b* (Fig. 2a). Upon linearization, we find that these interactions are at most of second order and

**FIG. 2.**
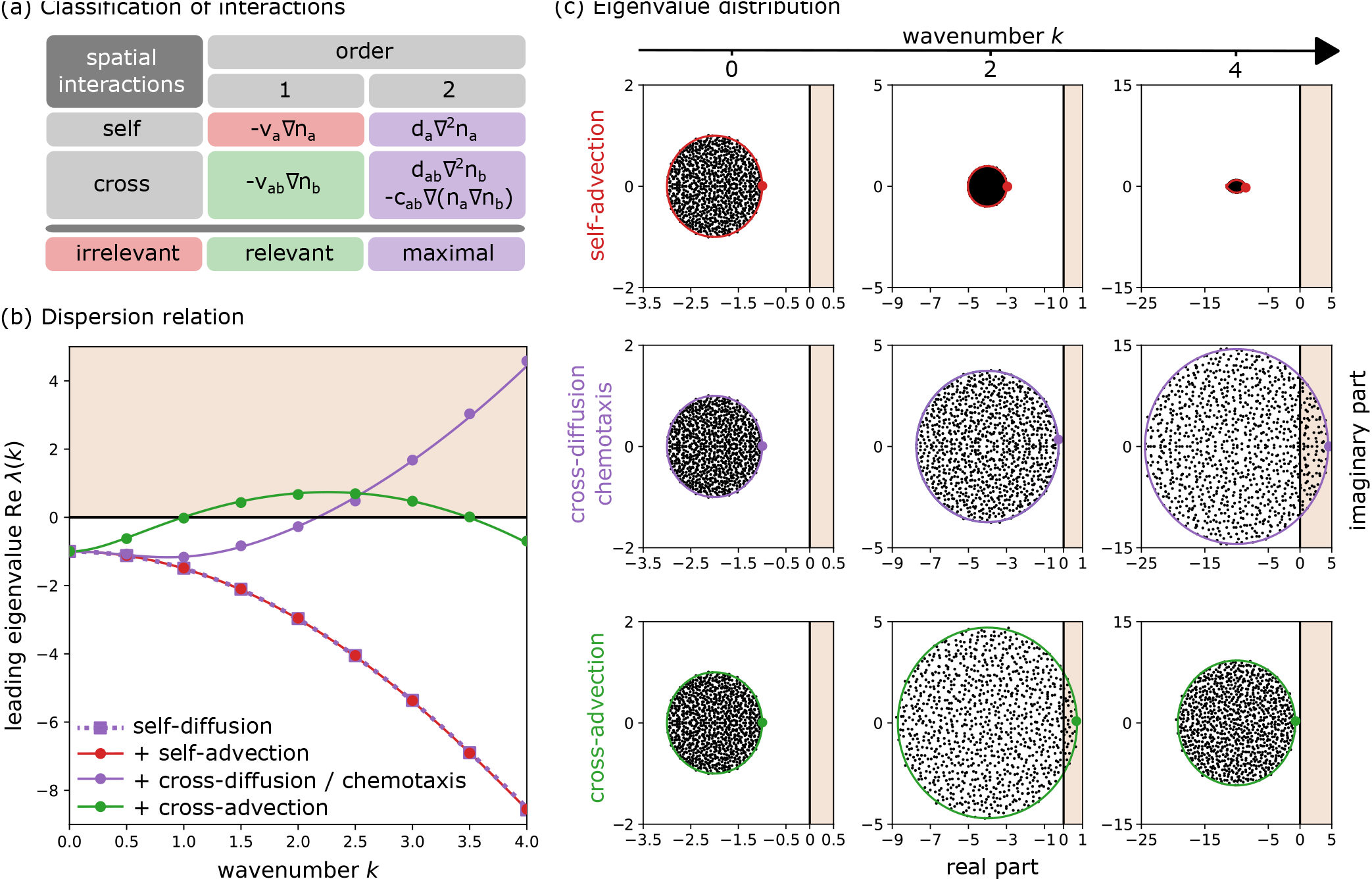
Example 1: Reaction-advection-diffusion systems. **(a)** Interactions in equation (13) are classified as irrelevant (red; self-advection, *v*_*a*_), relevant (green; cross-advection, *v*_*ab*_), and maximal (purple; self-diffusion, *d*_*a*_ *>* 0; cross-diffusion, *d*_*ab*_; chemotaxis, *c*_*ab*_). A positive self-diffusion (*d*_*a*_ *>* 0) is included in all our systems. **(b)** Dispersion relations for large (*N* = 1000) and unstructured systems (*τ*_*p*_ = 0) with different spatial interactions: self-diffusion alone (purple squares and dotted line), self-diffusion and self-advection (red dots and line), self-diffusion and cross-diffusion/chemotaxis (purple dots and solid line), self-diffusion and cross-advection (green dots and line). Lines correspond to the analytics in equation (A7) (resp. (B2)) if cross-interactions are present (resp. absent). Symbols represent numerical simulations of the Jacobian matrix in equation (14), where entries are sampled independently from Gaussian distributions (resp. a Gamma distribution for *d*_*a*_ *>* 0) with means *µ*_*s*0_ = −2, *µ*_*s*2_ = 0.5, *µ*_*s*1_ = *µ*_*cp*_ = 0 and standard deviations 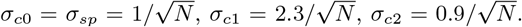. The brown region denotes a robust pattern formation. **(c)** Eigenvalues of the simulated Jacobian matrix for wavenumbers *k* = 0, 2, 4 and systems where self-diffusion is enhanced by self-advection (red), cross-diffusion/chemotaxis (purple) and cross-advection (green). Black dots represent all eigenvalues from the numerical simulations in panel (b), coloured dots represent the leading eigenvalues plotted in panel (b), and coloured lines represent the analytics, which are based on the same equations as in panel (b). See Supplemental Material for the Python code.

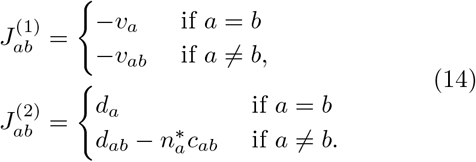

Since the temporal interactions are assumed to be unstructured, the spatially homogeneous stationary state is commonly approximated by 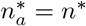 for some *n*^*\**^ [40, 42]. Therefore, by sampling the velocities, diffusivities and chemotactic coefficients independently and from identical distributions, we obtain equation (9) with

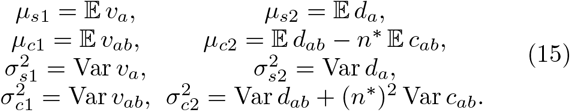

The models in mathematical biology commonly include a positive self-diffusion *d*_*a*_ *>* 0 [16], which implies that *µ*_*s*2_ *>* 0 and the maximal even interaction has order *P* = 2. In particular, as long as cross-advection is not anti-symmetric, the interactions can be classified as in Fig. 2a. By Theorem 2, a system with positive self-diffusion does not give rise to robust pattern formation (Fig. 2b). This result explains why self-diffusion alone cannot generate Turing patterns in large and unstructured random systems, as observed in numerical simulations [23, 31], and it formalizes the previous heuristic argument [33] and extends the previous analytics for homogeneous diffusivity with Var *d*_*a*_ = 0 [43]. In line with Theorem 2, we can also notice that the addition of self-advection into such a system cannot rescue the robustness of pattern formation (Fig. 2b,c). While self-advection cannot promote the robustness of pattern formation, Theorem 1 shows that sufficiently strong cross-diffusion and chemotaxis can generate patterns that are robust but not regular (Fig. 2b,c, Appendix D). This result aligns with the observation that patterns form robustly in the Keller-Segel model of chemotaxis [35], but these patterns blow up in finite time [44, 45]. Finally, Theorem 1 asserts that the only considered spatial interaction that gives rise to the formation of robust and regular patterns is cross-advection (Fig. 2b,c, Appendix D), aligning with the observation that cross-advection can support a variety of spatio-temporal patterns [46]. Overall, these results are consistent with a recent analysis of pattern formation in general reaction-advection-diffusion systems with two variables (*N* = 2), which suggests that cross-interactions facilitate pattern formation more robustly than self-interactions [47].

While the examples from mathematical biology are often limited to spatial interactions of order *p ≤* 2, higher order interactions appear commonly in soft condensed matter, such as fluid mixtures. In this context, spontaneous pattern formation, known as spinodal decomposition, corresponds to a phase transition in which a homogeneously distributed matter spontaneously separates into distinct phases [12]. Due to the inherent complexity of many physical systems [48], multiple studies have examined spinodal decomposition in systems with many components *N* [48–53]. For example, [52] studied a fluid mixture with *a* = 1, …, *N* components. The components were described by volume fractions *n*_*a*_(*t, x*), the remaining volume fraction of 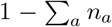 was attributed to the solvent, and the mechanics was described by the free energy functional

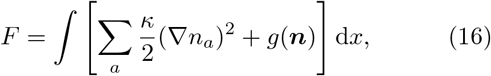

where *κ >* 0 describes a surface tension and

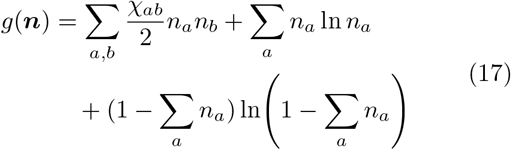

describes the bulk free energy density of the homogeneous state. In equation (17), the symmetric matrix *χ*_*ab*_ describes interactions between different components of the mixture and the remaining terms describe the entropy of the mixing. Spinodal decomposition has been examined using Landau-Ginzburg and random matrix theories [52]. Specifically, random matrix theory was used to test whether the Hessian matrix

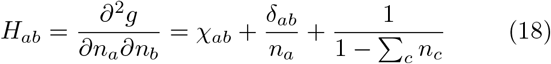

has a negative eigenvalue, that is, whether an appropriate homogeneous perturbation around a homogeneous stationary state 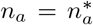 can reduce the free energy (Fig. 3a). Assuming that the symmetric interactions χ_*ab*_ are independent and identically distributed

**FIG. 3.**
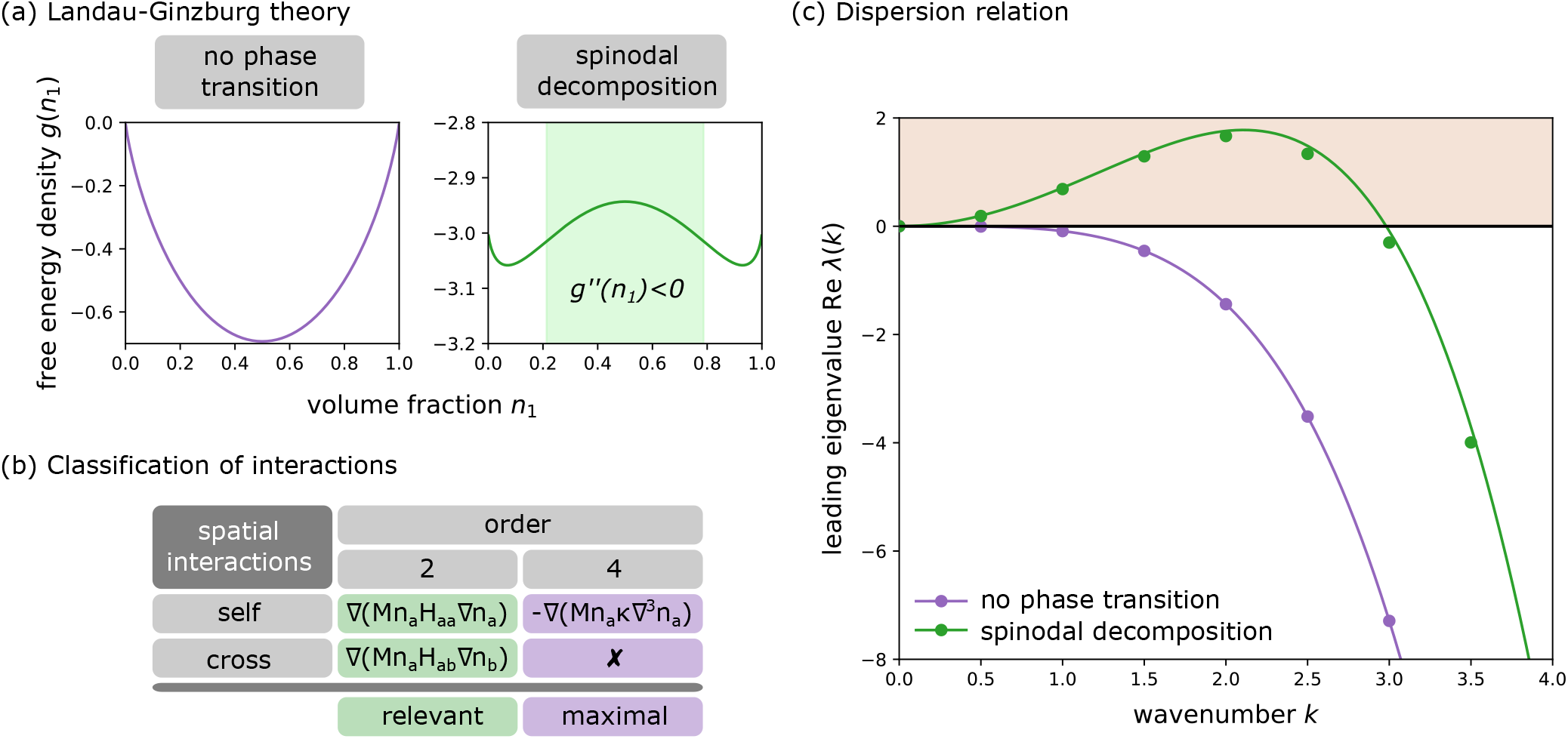
Example 2: Spinodal decomposition. **(a)** Landau-Ginzburg theory predicts whether a well-mixed fluid mixture spontaneously decomposes (spinodal decomposition) based on the free energy density *g*(***n***) in equation (17), depicted for a binary mixture (*N* = 2) with *n*_1_ + *n*_2_ = 1 and interactions 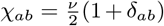 (*ν* = 0 left, *ν* = 3 right). Spinodal decomposition is expected to occur if the Hessian matrix has a negative eigenvalue (i.e. *g*^′′^(*n*_1_) < 0, light green region around the maximum of the right panel). **(b)** Interactions in equation (21) are classified as relevant (green; interactions between mixture components) and maximal (purple; surface tension *κ*). **(c)** Dispersion relations for a multi-component fluid mixture (*N* = 1000) around an equimolar homogeneous stationary state 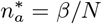 with *β* = 0.9 in the absence (purple dots and line) and presence (green dots and line) of unstructured symmetric interactions between different components (*τ*_*p*_ = 1). Lines plot the analytics in equation (A7). Symbols represent numerical simulations of the Jacobian matrix in equations (18), (19) and (23), where entries are sampled independently from a Gaussian distribution with a mean *ν*_*c*_ = *ν*_*s*_ = −6 and a standard deviation 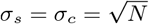, and the surface tension is chosen to be *κ* = *N/*10. The brown region denotes robust pattern formation. See Supplemental Material for the Python code.

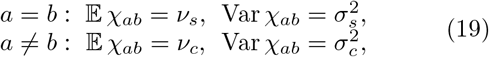

and that the homogeneous stationary state is equimolar (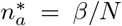 for some *β* < 1), spinodal decomposition occurs precisely when

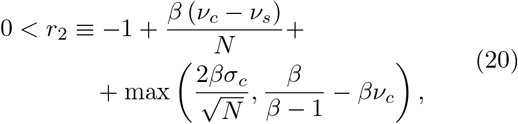

which coincides with the result in [52] for the restricted case of *ν*_*s*_ = *ν*_*c*_ = *ν* and *σ*_*s*2_ = *σ*_*c*2_ = *σ* (Appendix E). To simulate spinodal decomposition numerically, [52] used the so-called model B, given by

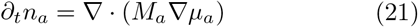

Where

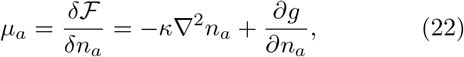

is the chemical potential of component *a* and *M*_*a*_ = *Mn*_*a*_ with *M >* 0 specifies the mobility of component *a*. Our method allows us to explore the dynamics in (21) analytically by observing that the growth of perturbations with wavenumber *k* around the homogeneous stationary state 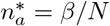 is governed by the Jacobian in equation (6), given by

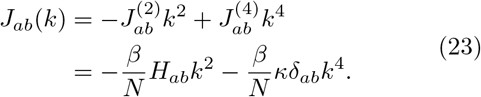

Using the same approach as in the proof of Theorem 1, it can be deduced that the real part of the leading eigenvalue *λ*(*k*) is given by

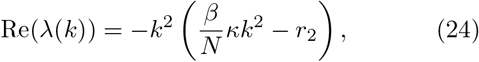

where *r*_2_ is defined by equation (20) and coincides with the definition in equation (11) (Appendix E). Therefore, in addition to learning that spinodal decomposition occurs when *r*_2_ *>* 0, we also learn that perturbations grow when their wavenumber is in the range of

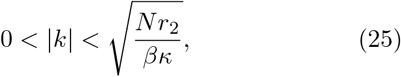

and the growth is maximal at the wavenumber

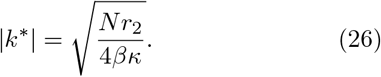

Not only does our framework allow for dynamical predictions beyond the standard Landau-Ginzburg theory, but it also provides a simple rule for determining which terms of the free energy functional in equation (16) robustly promote spinodal decomposition. Noticing that the dynamics in equation (21) increases the order of spatial interactions by +2 due to two spatial derivatives, we can classify the spatial interactions straight from the free energy functional in equation (16) as in Fig. 3b. Since the interaction strength of the maximal interaction *r*_4_ = −*βκ/N* is negative, Theorem 1 implies that the maximal interaction generated by surface tension cannot in itself generate spinodal decomposition but rather ensures that the spinodal decomposition is regular when generated by other terms (Fig. 3c). Moreover, Theorem 1 also implies that spinodal decomposition occurs when the interaction strength is sufficiently strong (*r*_2_ *≫* 0, Fig. 3c), which aligns with the more nuanced result in equation (20). Furthermore, if we generalized the surface tension *κ >* 0 to a positive definite matrix *κ*_*ab*_ via

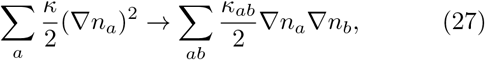

then Theorem 1 would imply that the newly introduced maximal cross-interactions of order *p* = 4 with negative strength *r*_4_ < 0 would still regularize the spinodal decomposition rather than promote its robustness. Similarly, the addition of symmetric spatial interactions of the form *n*_*a*_ *∇ n*_*b*_ into the free energy functional in equation (16) would introduce irrelevant interactions of order *p* = 3, meaning that such interactions do not improve the robustness of spinodal decomposition.

On the one hand, Theorem 1 provides a useful guide for constructing models that robustly admit spontaneous pattern formation by identifying appropriate spatial interactions order by order. On the other hand, some spatial interactions inherently carry a multitude of different orders *p*. Most notably, these interactions include non-local mechanisms, such as cell-cell interactions via pseudopodia in the context of developmental biology [54, 55] or long-range seed dispersal in the context of plant ecology [26, 56]. For example, the models for non-local interactions in [38, 57] have the general form

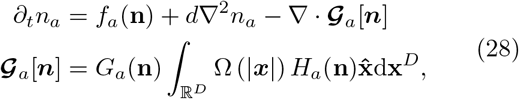

where *d >* 0 is diffusivity, *G*_*a*_(**n**) and *H*_*a*_(**n**) modify the interaction strength, Ω(***x***) is the interaction kernel and 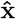 is the unit vector of a specified direction in the *D*-dimensional space ℝ^*D*^. As proved in [38], the Jacobian matrix of equation (6) is given by

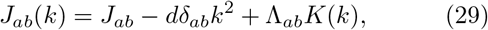

where *k* = | ***k***| is the magnitude of the wavenumber vector, the strength of the non-local interactions is captured by the matrix

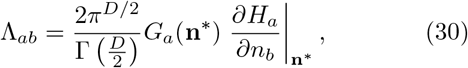

and *K*(*k*) is the transformed interaction kernel

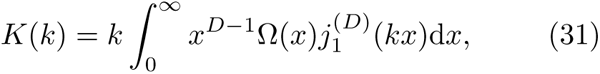

with 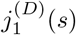 being the *D*-dimensional hyperspherical Bessel function. Since these hyperspherical Bessel functions are odd and analytic, the function *K*(*k*) is even and analytic, implying the corresponding term of the Jacobian in equation (29) includes interactions of infinitely many even orders *p*. For example, if Ω(*x*) = *e*^−*x*^ and two spatial dimensions are considered (*D* = 2), then the transformed interaction kernel is [38]

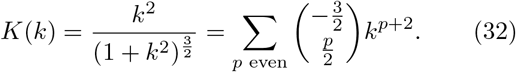

If we used Theorem 1 heuristically, this observation would imply that non-local interactions include infinitely many relevant interactions and pattern formation is robust (Fig. 4a). While these interactions are not independent, as assumed in Theorem 1, and are related to the same matrix Λ_*ab*_, the steps in the proof of Theorem 1 (Appendix A) can still be used. When the interactions Λ_*ab*_ are independent and identically distributed with

**FIG. 4.**
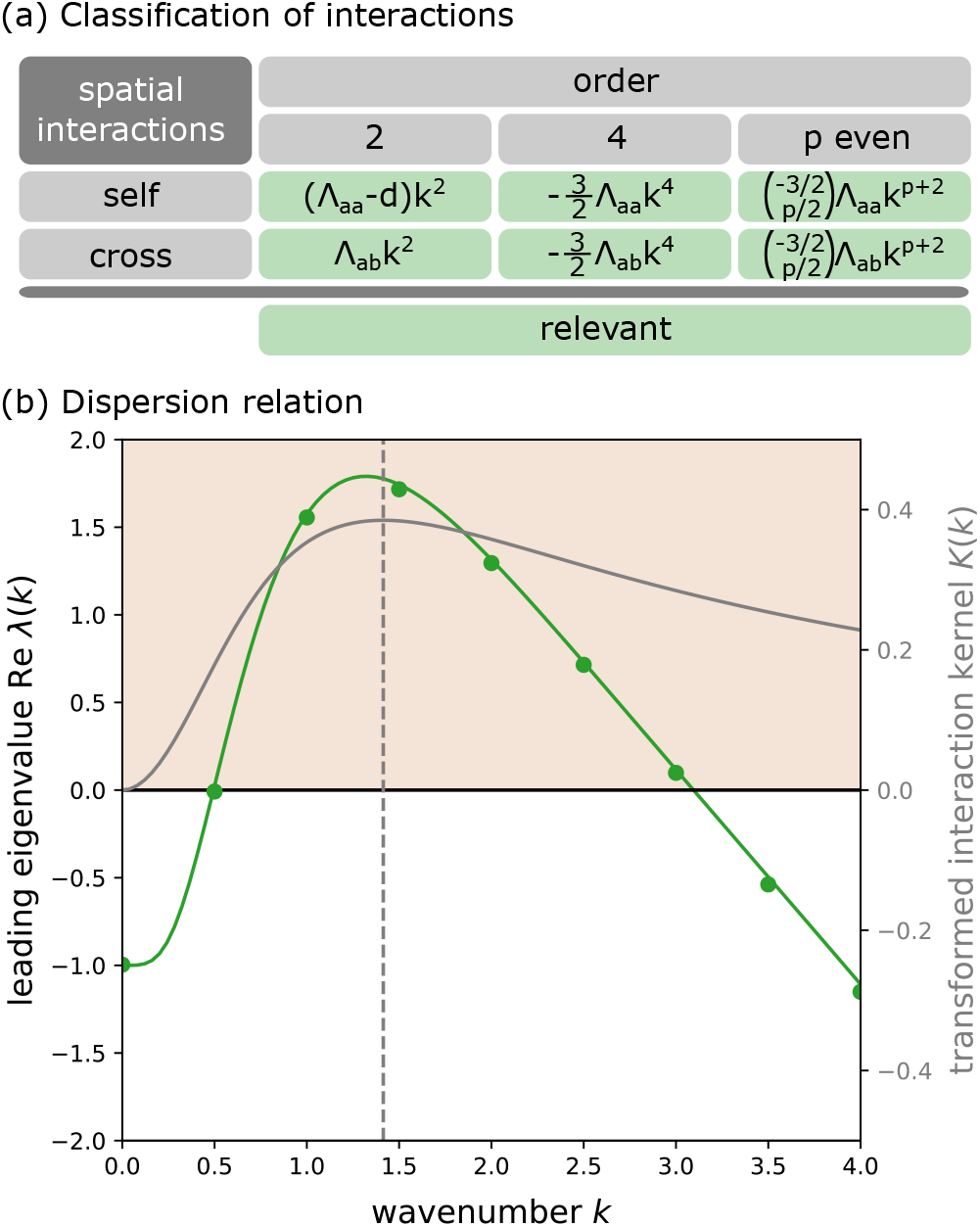
Example 3: Nonlocal interactions. **(a)** Nonlocal interactions in equation (29) are all relevant when the transformed interaction kernel is expanded in the powers of the wavenumber *k*. The interaction kernel *K*(*k*) in equation (32) is considered for illustration. **(b)** Dispersion relations for non-local interactions in a large system (*N* = 1000) with unstructured interactions (*τ*_*p*_ = 0). The green line corresponds to the analytics in equation (34). The green dots represent numerical simulations of the Jacobian matrix in equation (29) and (33), where entries are sampled independently from Gaussian distributions with means *µ*_*s*0_ = 2, *µ*_*c*0_ = *µ* =0 and standard deviations 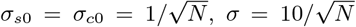 and the diffusion is chosen to be *d* = 0.1. The brown region denotes robust pattern formation. See Supplemental Material for the Python code. The dispersion relation is overlaid with the transformed interaction kernel *K*(*k*) in equation (32) (solid gray line) and its global maximum at 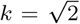 (dashed gray line), which approximately matches the maximum in the dispersion relation.

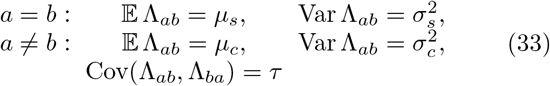

and the number of interacting variables *N* is large, the leading eigenvalue *λ*(*k*) of the Jacobian matrix in equation (29) is given by

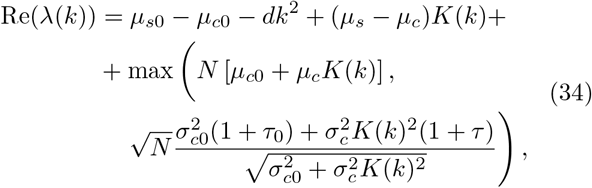

as proved in Appendix F. Assuming that *K*(*k*) *→ ∞* 0 as *k → ∞* (which is true for the commonly studied interaction kernels [38]), the value of Re(*λ*(*k*)) remains bounded from above by 0 as *k → ∞* and any spontaneous pattern formation must be regular. Moreover, since there exists a wavenumber *k* and a sign *s* = *±*1 such that *sK*(*k*) *>* 0, it is always possible to make the strength of non-local interactions

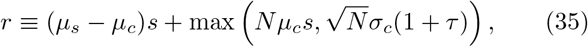

sufficiently large so that Re(*λ*(*k*)) is positive. Put differently, sufficiently strong non-local interactions can always robustly and regularly generate patterns (Fig. 4b), which aligns with our intuition based on the heuristic application of Theorem 1. Further analysis shows that the maximally growing perturbation has a wavenumber that can be approximated by the maximum of the transformed interaction kernel |*K*(*k*) | (Appendix F, Fig. 4b), further supporting the intuition that the non-locality of spatial interactions is the key ingredient for robust pattern formation in equation (28). These results generalize previous work that focused on pattern formation with multiplicative noise in the restricted case with *G*_*a*_ = *H*_*a*_ = *n*_*a*_ [57] and provides a key analytical insight into the improved robustness of pattern formation when non-local interactions are compared to reaction-advection-diffusion systems [38].

In this Letter, we have formalized the concept of robust pattern formation (Fig. 1) and illustrated this concept across a variety of models from mathematical biology (Fig. 2, 4) and soft condensed matter physics (Fig. 3). At a conceptual level, our approach shares similarities with the renormalization group in statistical and quantum field theories. While the renormalization group identifies field interactions that are relevant for the physics at large length scales, our method (Theorems 1 and 2) identifies interactions that are relevant for robust pattern formation. This approach allows one to formalize the observation that diffusion alone does not generate patterns robustly in large and unstructured systems [23, 31, 33] and that advection [17, 34], chemotaxis [35], cross-diffusion [36] and non-local interactions [37, 38] indeed facilitate pattern formation more robustly than diffusion (Fig. 2, 4). Moreover, our method can be used to explain the robustness of spinodal decomposition in soft condensed matter and analyze its dynamics beyond the regimes predicted from Landau-Ginzburg theory (Fig. 3). Our new tool can also offer a simple guide for creating mathematical models that explain pattern formation in highly complex physical systems [3–6, 21, 22]. Since physical systems can be potentially described by a large number of variables *N*, the systems that form patterns must either include relevant spatial interactions as outlined in Theorem 1 or have highly correlated and structured interactions. For example, if diffusion is the only mechanism at play, patterns can form in systems with a large number of variables *N* only if these systems are structured into activator and inhibitor subsystems [29, 58, 59], which might explain the complications with observing Turing patterns in real physical systems [28]. When a complex empirical system forms patterns and there is no apparent interaction structure, Theorem 1 lists spatial interactions that could robustly generate these patterns and can provide a guide for further empirical tests. Altogether, our work defined and examined the robustness of pattern formation in realistically complex systems, providing a map for searching specific mechanisms that generate these spatial patterns.

This work was supported by Oxford Mathematical Institute Scholarship (V.P.). For the purpose of Open Access, the author has applied a CC BY public copyright licence to any Author Accepted Manuscript (AAM) version arising from this submission.

## Appendix A: Proof of Theorem 1

Equations (6) and (9) imply that the on-diagonal and off-diagonal entries of the matrix *J*_*ab*_(*k*) are independent and identically distributed with

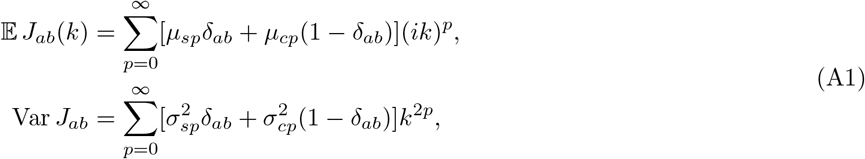

where the entries are independent up to a correlation between opposite entries across the diagonal, described by the pseudo-covariance

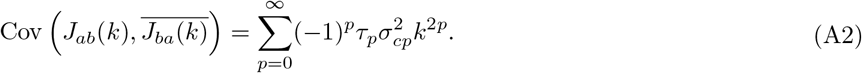

Since the off-diagonal entries generally have non-zero means, *J*_*ab*_(*k*) has a single outlier eigenvalue [60], given by

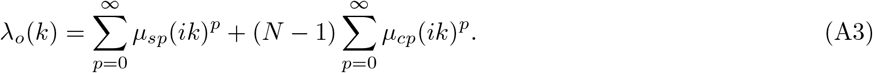

The distribution of the remaining eigenvalues can be determined by a rank-one perturbation approach, as in [40]. Specifically, the eigenvalues of the matrix *J*_*ab*_(*k*) have the same distribution as the eigenvalues of the matrix *J*_*ab*_(*k*) − 𝔼 *J*_*ab*_(*k*), up to a shift by

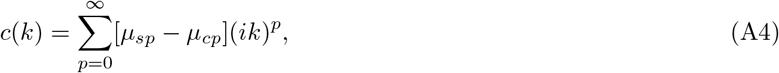

and the eigenvalues of the matrix *J*_*ab*_(*k*) − E *J*_*ab*_(*k*) are distributed uniformly in an ellipse [61] with a half-width *a*_+_(*k*) (resp. half-height *a*_−_(*k*)) given by

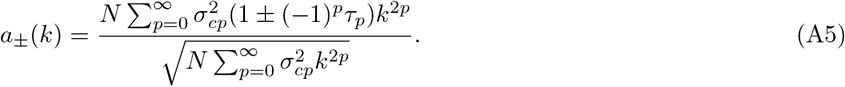

Therefore, the bulk of the eigenvalues of *J*_*ab*_(*k*) are distributed in an ellipse with center *c*(*k*), half-width *a*_+_(*k*) and half-height *a*_−_(*k*), implying that the eigenvalue in the bulk spectrum with the maximal real part is given by

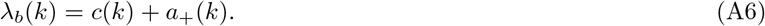

Thus, the eigenvalue with the maximal real part *λ*(*k*) is given by *λ*_*o*_(*k*) (resp. *λ*_*b*_(*k*)) if Re*λ*_*o*_(*k*) *>* Re*λ*_*b*_(*k*) (resp. Re*λ*_*o*_(*k*) < Re*λ*_*b*_(*k*)) and the corresponding maximal real part is

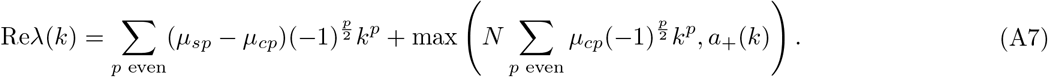

If the only interactions present had order *p*, then equation (A7) would imply that

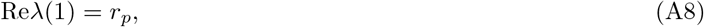

where *r*_*p*_ is the interaction strength of order *p* in equations (11) and (12) of Definition 1. Therefore, the sign of the interaction strength *r*_*p*_ in Definition 1 determines whether all eigenvalues of the matrix 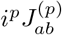 have negative real parts (*r*_*p*_ < 0) or not (*r*_*p*_ *>* 0). In particular, since we assume that the homogeneous stationary state is stable, it must be true that all eigenvalues of 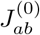 have negative real parts and *r*_0_ < 0. With these relationships, we can prove the respective statements in Theorem 1:

1. We can notice that the equation (A7) does not include *µ*_*sp*_, *µ*_*cp*_ and 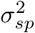 for any self-interaction of any odd order *p*. Moreover, for any odd order *p* with *τ*_*p*_ = 1, Re*λ*(*k*) decreases with increasing 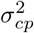, as can be deduced from equation (A5). Therefore, pattern formation is not robust when irrelevant interactions are the only interactions present since

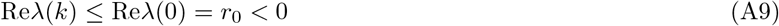

for any wavenumber *k*.
2. When there is at least one significant interaction of order *p > P* with non-vanishing strength *r*_*p*_ ≠ 0, equations (A7) and (A5) imply that

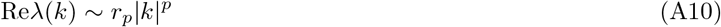

as |*k*|*→ ∞*, where *p* corresponds to the maximal order among all such significant interactions. Since *r*_*p*_ *≥* 0 by equation (12), the presence of significant interactions makes pattern formation robust and irregular.
3. When all significant interactions have vanishing strength and the relevant interactions are absent, there are two cases that depend on the strength of the maximal interaction *r*_*P*_, given by equation (11).
  a. When *r*_*P*_ *>* 0, the substitution of equations (11) and (A5) into (A7) imply that

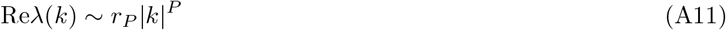
  b. as |*k*| *→ ∞* and pattern formation is robust and irregular.
  c. When *r*_*P*_ *≤* 0, equation (11) implies that

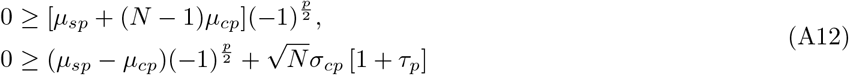

for *p* = *P*. Equation (A12) is also true for *p* = 0 (with strict inequalities) due to the stability of the homogeneous stationary state (*r*_0_ < 0). Therefore, using equation (A12) for *p* = 0, *P*, we find that equation (A3) for the outlier eigenvalue satisfies

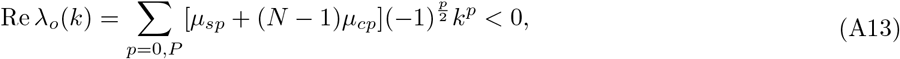

and equation (A6) for the eigenvalues in the bulk spectrum satisfies

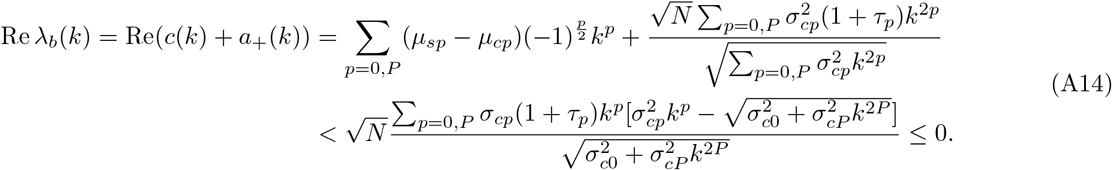

for any wavenumber *k*. Therefore, equations (A13) and (A14) imply that the maximal real part of any eigenvalue satisfies

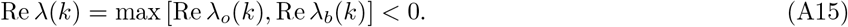

for any wavenumber *k*, meaning that pattern formation is not robust according to Definition 2.
4. If we fix some wavenumber *k*^*\**^ and some relevant interaction of order *p*, we can make the interaction strength sufficiently strong (*r*_*p*_ *≫* 0) so that

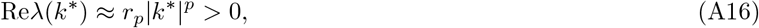

implying that pattern formation is robust for sufficiently strong relevant interactions. Moreover, when all significant interactions have vanishing strength and the maximal interaction has negative strength, equation (A11) is valid asymptotically at large *k* and pattern formation is regular.

## Appendix B: Proof of Theorem 2

To prove Theorem 2, we treat particular realizations 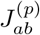 of the spatial interactions *p ≥* 1 as deterministic matrices and the matrix of temporal interactions 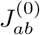 as a sum of a diagonal deterministic matrix and a non-diagonal random matrix, as in equation (9) with *τ*_0_ = *µ*_*c*0_ = 0. Assuming that the only spatial interactions are self-interactions (i.e., 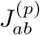 are diagonal matrices), the Jacobian matrix in equation (6) can be written as

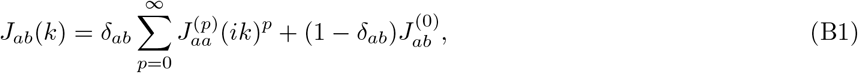

where the first term corresponds to a deterministic diagonal matrix and the second term corresponds to an off-diagonal stochastic matrix. Using the result in [39], the eigenvalues of the Jacobian matrix *J*_*ab*_(*k*) are distributed within a region of the complex plane *z* ∈ ℂ, given by

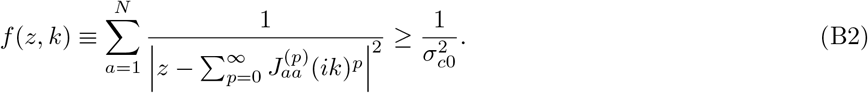

Since the temporal dynamics is assumed to be stable in the absence of spatial interactions, the region given by equation (B2) lies in the real negative half-plane {Re(*z*) < 0} when *k* = 0. Specifically, this implies that 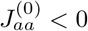 for all *a*. With these relationships, we can prove the respective statements in Theorem 2:

1. When the irrelevant interactions are the only interactions present, we can use the inequality | *z*| *≥* | Re(*z*) | to note that

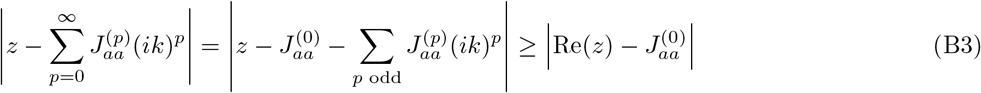

for every field variable with index *a*. Therefore,

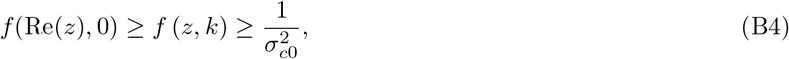

meaning that, if *z* lies in the eigenvalue region given by equation (B2) for some *k* ≠ 0, then Re(*z*) lies in this eigenvalue region for *k* = 0. Since all eigenvalues have negative real parts at *k* = 0, this observation implies that all eigenvalues have negative real parts at all wavenumbers *k*. Put differently, pattern formation is not robust.
2. When all relevant interactions are absent, there are two cases depending on the signs of the maximal interactions 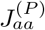, where we recall that *P* is even by Definition 1.
  a. If 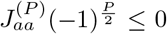 for all *a*, we start by considering a case where all irrelevant interactions are absent. In this case, the function *f* (*z, k*^2^) in equation (B2) only includes the power *k*^*P*^ and

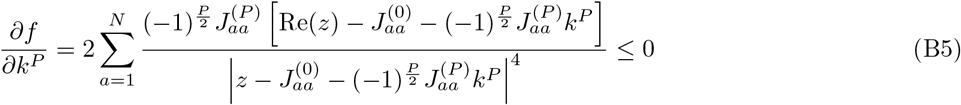

for any *z* with a positive real part, since 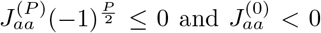 by assumption. This observation implies that, for any wavenumber *k*,

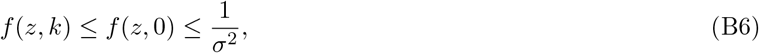

where the second inequality follows from the fact that the eigenvalues have negative real parts at *k* = 0. Therefore, the eigenvalue region in equation (B2) lies in the negative real half plane for any wavenumber *k* and pattern formation is not robust. To extend this result to cases where irrelevant interactions are present, we can prove that a system with irrelevant interactions has at least one eigenvalue with a positive real part only if there is an eigenvalue with a positive real part in the system that has the irrelevant interactions removed. To prove this statement, we can use the inequality |*z*| *≥* | Re(*z*)| so that

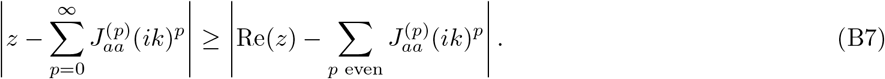 Defining

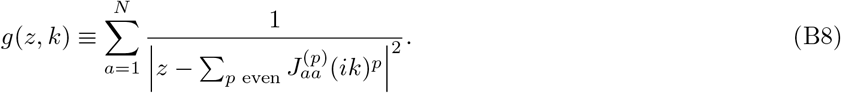

to be the function as in equation (B2)but for a system where the irrelevant interactions are removed, equation (B7) implies that

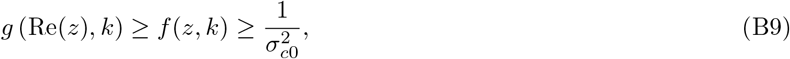

where *f* (*z, k*) is still given by equation (B2) and describes the system where irrelevant interactions are present. Therefore, if the eigenvalue region in equation (B2) for the system with irrelevant interactions (described by function *f*) contains a number *z* with a positive real part, then the positive number Re(*z*) is included in the eigenvalue region of the system that has the irrelevant interactions removed (described by function *g*). But, as we have just shown, such a system only has eigenvalues with negative real parts. Therefore the statement (a) of Theorem 2 applies even if irrelevant interactions are present.
  b. When 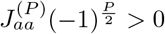 for some *a*, then the region given by equation (B2) includes the circle

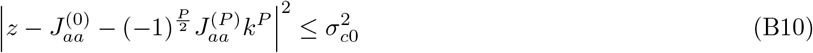

centred at

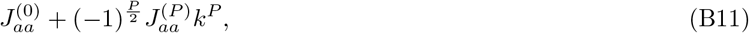

which is positive for any sufficiently large *k* because 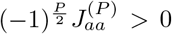 by assumption and *P* is an even power. Therefore, in this case, pattern formation is robust and irregular.
3. Since the eigenvalue region in equation (B2) includes the circles for each field variable of index *a*

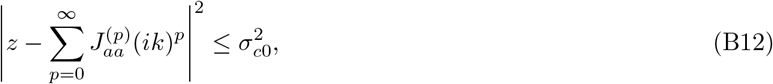

with the real part of the centre located at

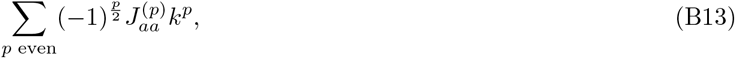

sufficiently strong relevant interactions (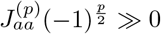 for some *a*) can always move this centre to the positive real half plane at any fixed wavenumber *k*. Therefore, sufficiently strong relevant interactions can always make pattern formation robust as in Definition 2. Moreover, for sufficiently large *k*,

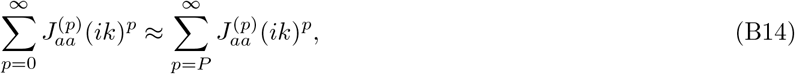

and the eigenvalue region in equation (B2) can be approximated by the corresponding eigenvalue region with interactions of order *p* < *P* removed, which includes all relevant interactions. When 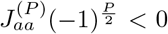 for all *a*, the second point of Theorem 2 implies that such a reduced system has only eigenvalues with negative real parts, meaning that all eigenvalues of the full system must have negative real parts at sufficiently large wavenumbers *k*. Therefore, the resulting pattern formation is robust when 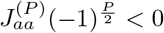 for all *a*.

## Appendix C: Extension to Multiple Spatial Dimensions

In the case of multiple spatial dimensions, the situation is complicated by the fact that the interactions 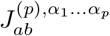 in equation (6) now have multiple upper spatial indices. Writing ***α*** = (*α*_1_, …, *α*_*p*_), the independent and identical distribution of interactions in equation (9) can be generalized to

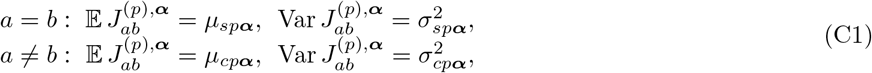

and equation (10) to

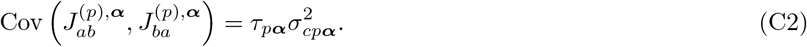

Correspondingly, equations (A1) and (A2) are generalized to

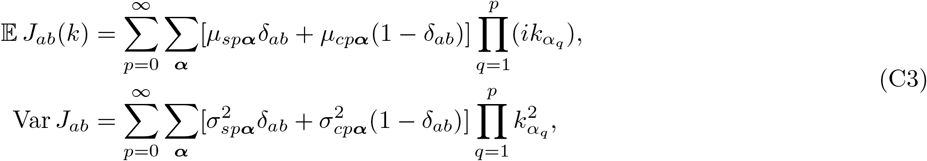

And

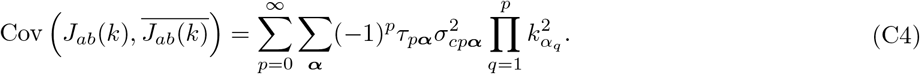

With these modifications, one could appropriately modify the subsequent steps in the proof of Theorem 1 (Appendix A) and Theorem 2 (Appendix B) to generalize these results to multiple spatial dimensions.

For illustration, we can notice that Theorem 1 generalizes to an arbitrary number of spatial dimensions *D* when the interactions are minimally anisotropic, that is, when

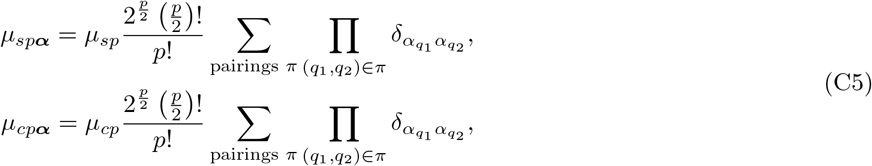

for interactions of even order *p* and

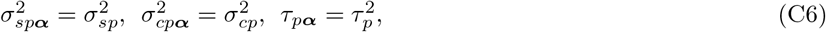

for interactions of all orders *p*. In this case, the key equation (A7) in the proof of Theorem 1 remains valid if we interpret *k* = |***k***| as the magnitude of the wavenumber vector ***k***. Therefore, Theorem 1 remains valid in arbitrary spatial dimension *D* under these conditions.

## Appendix D: Reaction-Advection-Diffusion Systems

In this section, we analyze the reaction-advection-diffusion system in equations (13), (14) and (15), with the additional assumption that cross-interactions are not constrained by any symmetries, that is *τ*_*p*_ = 0. Under this assumption, the eigenvalue with the maximal real part in equation (A7) becomes

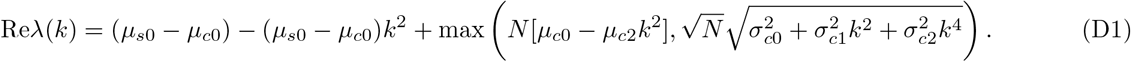

When *k* is large, equation (11) implies that

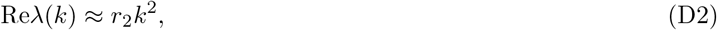

which means that pattern formation is robust and irregular when

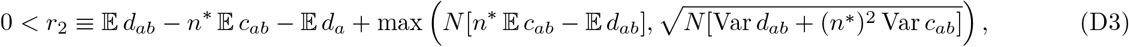

that is, when chemotaxis is sufficiently positive (E *c*_*ab*_ *≫* 0), or cross-diffusion sufficiently negative (𝔼 *d*_*ab*_ *≪* 0) or either of these is sufficiently heterogeneous (Var *d*_*ab*_, Var *c*_*ab*_ *≫* 0). When *r*_2_ < 0, the stability of the homogeneous stationary state implies *r*_0_ < 0 (as shown in Appendix A) and we obtain

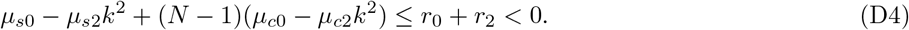

Therefore, pattern formation is robust precisely when

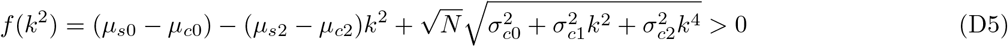

for some wavenumber *k*, which can be rewritten as

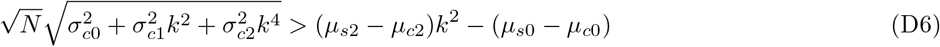

Since the interaction strengths *r*_0_ and *r*_2_ in equation (11) are assumed to be negative, it is true that

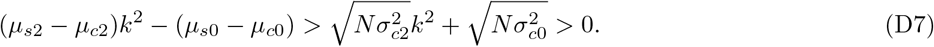

Upon squaring and rearranging the equation (D6), we learn that pattern formation is robust precisely if

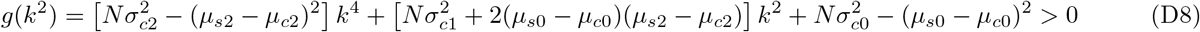

for some wavenumber *k*. Since the interaction strengths *r*_0_ and *r*_2_ in equation (11) are assumed to be negative, the coefficients at *k*^4^ and *k*^0^ are negative, meaning that the resulting pattern formation is necessarily regular. To determine when this robust and regular pattern formation occurs, we can notice that the function *g*(*k*^2^) has a maximum at

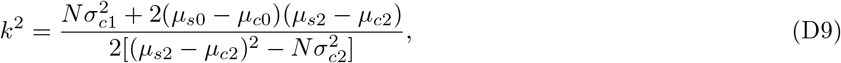

Provided 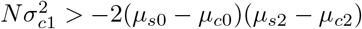, and *f* (*k*^2^) is positive at this maximum, provided

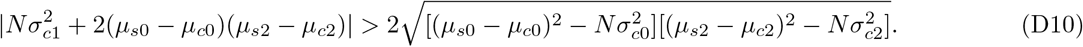

Therefore, there is an intermediate range of wavenumbers *k* corresponding to growing perturbation modes precisely if

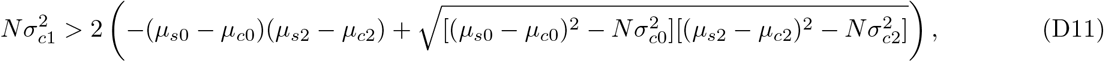

that is, precisely when the cross-advection is sufficiently heterogeneous (i.e., 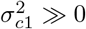). When this condition is satisfied, pattern formation is robust, regular and the maximally growing perturbations have wavenumbers given by equation

(D9). In particular, we reproduce the results of Theorem 1 by showing that patterns can be generated robustly by sufficiently strong cross-diffusion, chemotaxis and cross-advection, out of which only cross-advection generates patterns regularly. By performing this more detailed calculation, we additionally obtain the exact conditions for robust pattern formation that is irregular (equation (D3)), and regular (equation (D11)), as well as the critical wavenumber of the fastest growing perturbation for the latter case (equation (D9)).

## Appendix E: Spinodal Decomposition of Soft Condensed Matter

In this section, we analyze the spinodal decomposition as described by equations (18), (19) and (23). Upon substituting equation (18) into equation (23) and applying equation (19), we can see that

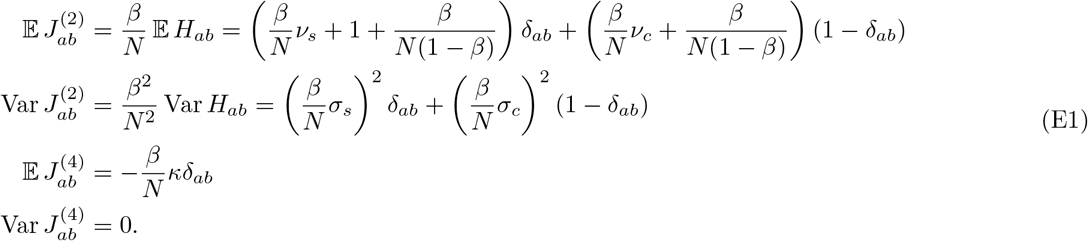

real part *λ*(*k*) must satisfy

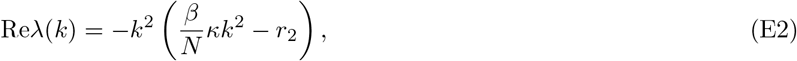

where *r*_2_ is given by equation (20). Substitution of these relationships into equation (11) also reveals that *r*_2_ in equation (20) coincides with the interaction strength in Definition 1. Since Re*λ*(*k*) is a quadratic function, it is straightforward to deduce that patterns form robustly precisely when *r*_2_ *>* 0. Moreover, such pattern formation is always regular and restricted to wavenumbers in the range of equation (25), with the fastest growing perturbation having a critical wavenumber given by equation (26). Finally, to relate our dynamical approach with the approach of Landau-Ginzburg theory, we note that *r*_2_ *>* 0 precisely when the matrix 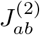 has at least one positive eigenvalue (see Appendix A) which, by equation (23), happens precisely when the matrix *H*_*ab*_ has at least one negative eigenvalue.

## Appendix F: Nonlocal Interactions

In this section, we analyze the model with non-local interactions, as described in equations (29) and (33). Using these equations, we can deduce that

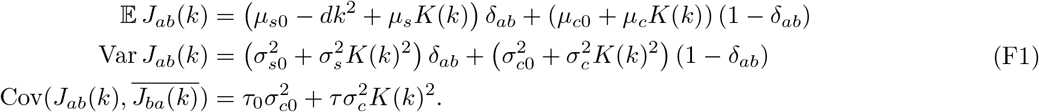

Assuming that the variance of the diagonal elements is bounded by the variance of the off-diagonal elements as in Theorem 1, we can use the same arguments from random matrix theory as in Appendix A [40, 60, 61] to deduce that the Jacobian matrix *J*_*ab*_(*k*) has a single outlier eigenvalue

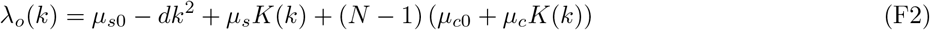

and a bulk spectrum of eigenvalues, with a point of maximal positive real part that is given by

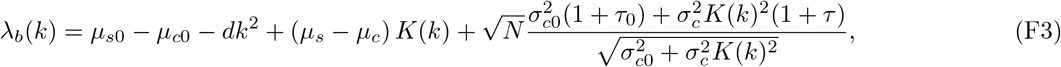

for a fixed wavenumber *k*. Noticing that these eigenvalues are real, we can deduce that the eigenvalue with the maximal real part is given by

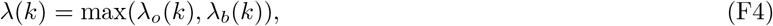

which, upon substitution of equations (F2) and (F3), proves equation (34) from the main text. Defining

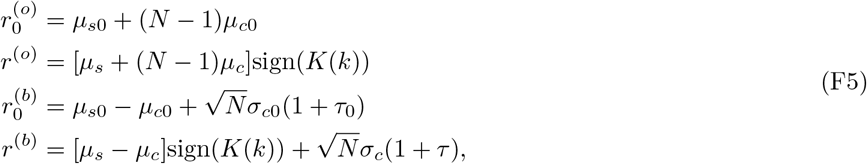

we deduce that the interaction strengths in equations (11) and (20) are given by

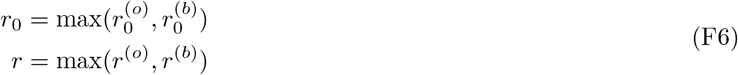

and the eigenvalues in equations (F2) and (F3) are given by

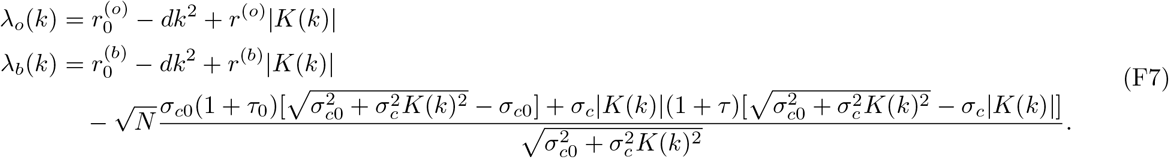

Since the homogeneous stationary state is stable by assumption, we know that *r*_0_ < 0, meaning that 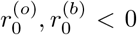. Therefore, the leading eigenvalue *λ*(*k*) is positive precisely when either of *λ*_*o*_(*k*) and *λ*_*b*_(*k*) is positive, which happens when *r*^(*o*)^ (resp. *r*^(*b*)^) is sufficiently large. This argument implies that sufficiently strong non-local interactions with *r ≫* 0 can robustly generate patterns, as presented in the main text. Moreover, when the strength of the non-local interaction *r*^(*i*)^ *≫* 0 is sufficiently strong for *i* = *o* (resp. *i* = *b*), the corresponding eigenvalue in equation (F7) can be approximated by

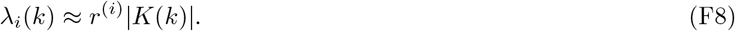

This observation implies that, when patterns form due to the bulk *i* = *b* (resp. outlier *i* = *o*) eigenvalue being positive (i.e., when *r*^(*i*)^ *≫* 0), the critical wavenumber *k* that maximizes *λ*_*i*_(*k*) is given by the maximum of |*K*(*k*) |. Therefore, when patterns form, the critical wavenumber *k* that maximizes the leading eigenvalue *λ*(*k*) can be approximated by the maximum of |*K*(*k*)|, as stated in the main text.

